# GREEN: a lightweight architecture using learnable wavelets and Riemannian geometry for biomarker exploration

**DOI:** 10.1101/2024.05.14.594142

**Authors:** Joseph Paillard, Jörg F. Hipp, Denis A. Engemann

## Abstract

Spectral analysis using wavelets is widely used for identifying biomarkers in EEG signals. At the same time, Riemannian geometry enabled theoretically grounded machine learning models with high performance for predicting biomedical outcomes from multichannel EEG recordings. However, these methods often rely on handcrafted rules and sequential optimization. In contrast, deep learning (DL) offers end-to-end trainable models that achieve state-of-the-art performance on various prediction tasks but lack interpretability and interoperability with established neuroscience concepts. We introduce GREEN (Gabor Riemann EEGNet), a lightweight neural network that integrates wavelet transforms and Riemannian geometry for processing raw EEG data. Benchmarking on five prediction tasks (age, sex, eyes-closed detection, dementia diagnosis, EEG pathology) across three datasets (TUAB, CAUEEG, TDBRAIN) with over 5000 participants, GREEN outperformed non-deep state-of-the-art models and performed favorably against large DL models on the CAU benchmark using orders of magnitude fewer parameters. Computational experiments showed that GREEN facilitates learning sparse representations without compromising performance. The modularity of GREEN allows for the computation of classical measures of phase synchrony, such as pairwise phase-locking values, which are found to convey information for dementia diagnosis. The learned wavelets can be interpreted as bandpass filters, enhancing explainability. We illustrate this with the Berger effect, demonstrating the modulation of 8-10 Hz power when closing the eyes. By integrating domain knowledge, GREEN achieves a desirable complexity-performance trade-off and learns interpretable EEG representations. The source code is publicly available.

## 1 Introduction

Electroencephalography (EEG) is well established for assessing large-scale cortical dynamics (Buzsáki and Mizuseki, 2014) and has proven useful for analysis of sleep (Dement and Kleitman, 1957), anesthetic monitoring (Rampil, 1998) or clinical examination of seizures (Jasper, 1949). These applications have in common that the phenomena of interest are characterized by high-amplitude signals that can be visually assessed by trained clinicians. Over the past decades, research in clinical and cognitive neurosciences has led to a wide range of experimental methods and data-analysis strategies that have allowed researchers to quantify patterns of brain activity that are hard to detect from raw EEG traces (Bekinschtein et al., 2009; Frohlich et al., 2019a; Hawellek et al., 2022; Hipp et al., 2011; Klimesch et al., 1998; Polich, 2007; Tallon-Baudry et al., 1996). Furthermore, a coevolution of instrumentation (Casson, 2019; Fiedler et al., 2022), curation of reusable datasets (Kim et al., 2023; Obeid and Picone, 2016; van Dijk et al., 2022) and innovation in signal processing and machine learning (ML) (Delorme and Makeig, 2004) unfolded within the field of EEG. In combination, these factors hold promise to extend the clinical and biotechnological applications of EEG by facilitating the discovery of novel brain-activity biomarkers. In many traditional EEG applications, the signal power at specific electrodes or the global average signal power have often been sufficient for monitoring and phenotyping patient groups based on changes in oscillatory brain activity (Benca et al., 1999; Dressler et al., 2004; Otto, 2008). The logarithm of the power is often preferred as it reflects universal scaling laws of brain structure and function (Buzsáki et al., 2013; Buzsáki and Mizuseki, 2014). The logarithm also facilitates comparisons between frequencies along the spectrum which, due to inverse proportionality between power and frequency—the 1*/f* scaling—, is dominated by slower brain waves (Bedard et al., 2006; Pritchard, 1992). When the objective is to build predictive models of complex cognitive processes, pathologies or other biomedical variables, modeling approaches that make more elaborate use of the multivariate complex EEG signal may substantially enhance the signal-to-noise ratio (Cohen, 2022; Nikulin et al., 2011). Examples include motor imagery in brain-computer interfaces (BCI) (Ramoser et al., 2000; Wolpaw et al., 1991), subtle signal alterations related to disorders of the central nervous system (CNS) (Shaw, 2002) or pharmaceutical treatments (Hirano and Uhlhaas, 2021; Wu et al., 2020).

A fundamental problem that needs to be faced in such applications arises from electromagnetic field spread. Due to volume conduction, electrical potentials are spread out across all electrodes such that signals measured on the scalp—in the electrode space—imply nonlinear distortions (Dähne et al., 2014; Mosher et al., 1999; Nunez and Srinivasan, 2006; Sabbagh et al., 2020), e.g. electrode-space power distributions do not stand in a linear relationship to the source-power distributions of interest. Source localization is a method for uncovering local and long-range neural synchronization by unmixing the EEG signal by solving biophysical inverse problems before computing features (Hipp et al., 2011, 2012; Kudo et al., 2024; Ranasinghe et al., 2022). However, this is not always practical as source localization is most accurate with individual anatomic magnetic resonance imaging (MRI) and additional interactive work for coregistration between electrodes and the individual brain model. Furthermore, in a distributed brain model, thousands of candidate dipole locations are used, which can add complexity and often requires the choice of anatomical atlases for dimensionality reduction (Destrieux et al., 2010; Khan et al., 2018; Tait et al., 2021).

In this context, machine learning has provided powerful tools for uncovering complex brain signals arising from multichannel EEG recordings. For instance, by learning spatial filters using the between-channel covariance matrix as input, the problem of volume conduction and source mixing can be tackled without explicitly relying on biophysical source models (Dähne et al., 2014; Koles et al., 1990; Wu et al., 2020). The same approach can be used for enhancing local oscillatory patterns and subtracting background noise (Nikulin et al., 2011). Alternatively, Riemannian geometry provides a theoretical framework to derive representations for covariance matrices which mitigate distortions due to volume conduction by projecting them to a Euclidean space while preserving distances, hence allowing the use of classical ML tools (Barachant et al., 2010, 2013). When preceded by frequency filtering, these features can be used with standard regularized linear techniques to define models for predicting from subtle brain activity patterns (Sabbagh et al., 2020).

More recently, the rich literature of deep learning (DL) for speech processing and computer vision has been repurposed for the analysis of EEG signals, leading to several potential benefits (LeCun et al., 2015; Schmidhuber, 2015; Szegedy et al., 2015). First, DL models allow for end-to-end computation, i.e., combining temporal filtering, spatial filtering and estimation of nonlinear functions into one optimization problem (Lawhern et al., 2018; Schirrmeister et al., 2017). Furthermore, optimizing the parameters of these models via stochastic optimization methods (Kingma and Ba, 2017; Robbins and Monro, 1951) makes it possible to learn from large datasets despite the significant memory footprint of the raw signal. Finally, the huge functional space covered by these architectures combined with their modular nature enables learning complex functions, which holds promise to facilitate the discovery of previously neglected signals.

Despite major methodological advances, DL applied to EEG remains to unlock breakthroughs in clinical applications or biomarker discovery. In the EEG context, we have identified several limitations of the applied DL literature that can hinder the attractiveness of DL methods for neuroscientists. First, the architecture choices are most of the time directly adopted from speech processing or computer vision applications (Kim et al., 2023; Şeker and Özerdem, 2024; Zhang et al., 2021) As a consequence, the complexity of models increased along with data requirements and computational cost while missing an intelligible link with established neuroscientific results and concepts such as capturing the oscillatory dynamics of neural populations in the cortex using band-pass filtering or measuring phase coupling (Buzsáki and Draguhn, 2004; Buzsáki and Mizuseki, 2014; Hipp et al., 2011). This makes it more difficult to understand how the functions learned by DL relate to the EEG landmarks and phenomena investigated in the neuroscience literature, hence, limiting the utility as a tool in applications.

Parallel research efforts have led to first successes at incorporating established EEG processing operations. For example, while still applying a canonical convolutional architecture design, Schirrmeister et al. (2017) proposed a minimal architecture that mimicked the operations of classical common-spatial-pattern filter bank models (Koles et al., 1990). In a recent study, real-valued wavelets were used instead of classical convolution kernels (Barmpas et al., 2023). Another line of research has studied Riemannian geometry in DL models (Carrara et al., 2024; Liu et al., 2024; Wilson et al., 2022; Zhang and Etemad, 2024). It is, however, noteworthy that several DL architectures explored for EEG focus on the power of the signal (Barmpas et al., 2023; Schirrmeister et al., 2017; Wilson et al., 2022). Such DL models do not explicitly compute phase-related features, hence, potentially missing the connection with the vast EEG literature on phase coupling (Lachaux et al., 1999; Varela et al., 2001). Finally, the lack of constraints in these architectures can provoke unsustainable computational scaling requiring massive datasets that are not yet available for EEG (Thompson et al., 2024, 2020). To support further exploration of the untapped potential of EEG for medical applications and biomarker development through deep learning, it will, therefore, be important to integrate neuroscientific and biophysical prior knowledge into the architecture design of neural networks.

In this work, we make an attempt to address this challenge. We propose a new light-weight architecture termed *GREEN*, combining Gabor wavelets and Riemannian computation in a neural network for EEG data. The article is organized as follows. We first present a background review of the related literature, motivate the architecture choices and detail the different architectures studied in this work. We then present empirical benchmarks of model performance and complexity on three international datasets with EEGs from more than 5000 human participants in five prediction tasks. These benchmarks focus on studying improvement over robust and state-of-the-art baseline models based on Riemannian geometry as well as positioning our work against DL architectures. Finally, we demonstrate the utility of *GREEN* for exploring model complexity in terms filter bank sizes and learned representations in different prediction tasks.

## 2 Related work

Published DL architectures for EEG focus on end-to-end processing of raw EEG data (Kim et al., 2023; Roy et al., 2019; Schirrmeister et al., 2017) and provide state-of-the-art prediction performance on a range of tasks. However, the scaling in terms of the number of parameters of such models seems unsustainable given the scarcity of EEG data and their lack of interpretability hinders their impact for biomarker discovery (Thompson et al., 2020). In this background review, we focus on classical spectral analysis tools and Riemannian geometry to motivate architecture design choices for our work that aims at overcoming limitations of current DL applications to EEG.

**Spectral analysis** of EEG recordings is a key step for identifying clinically relevant features. However, the classical Fourier transform is not always well suited for such data since it lacks temporal resolution. Time-resolved methods such as the complex wavelet transform are often preferred as they not only capture temporal variations of the frequency content but also allow for frequency-dependent spectral smoothing. This work will focus on the wavelets introduced by Dennis Gabor, which present an optimal trade-off between time and frequency localization (Gabor and D., 1946). A Gabor wavelet *φ_f_*, consist of a complex sinusoid of frequency *f* modulated by a Gaussian window characterized by its standard deviation *σ_t_*,

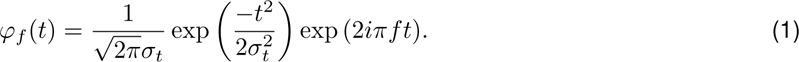

As many natural processes, the power of EEG signals have a 1*/f* scaling (Buzsáki and Mizuseki, 2014). To analyze such signals, wavelet families are constructed using the logarithmic frequency scale in octaves introduced by Jean Morlet (Morlet et al., 1982). This provides a non-orthogonal basis of wavelets that supports a wide array of techniques for studying brain function and can easily be interpreted as bandpass filters. In such parametrizations, wavelets are more localized in time and more spread in the frequency domain at high frequencies. This parametrization is therefore helpful for comparing brain activity along the frequency spectrum by providing physiologically appropriate smoothing (*σ_t_* log-linearly decreasing with *f*) which facilitates visualization and biomarker exploration (Frohlich et al., 2019a,b; Hawellek et al., 2022; Hipp et al., 2011; Tallon-Baudry et al., 1996).

In practice, neuroscientists use large filter banks to capture the entire frequency content of the signal. From a theoretical perspective, this does not lead to *optimal wavelet bases* in the sense of a minimally sized complete representation of the signal (Mallat, 1999) and, as a result, Morlet filter bank features are often non-sparse and highly correlated. Combining such Morlet-parametrized wavelets and ridge regularization (Hoerl and Kennard, 1970) has proven effective in a recent study where the authors observed that prediction performance saturated at a certain filter bank size but did not degrade (Bomatter et al., 2023).

While proving that this approach works well for regularized linear models, such non-sparse representations can lead to intractable computation costs for complex model architectures. To mitigate this issue, the parameters of the wavelet can be learned in a neural network in order to replace handcrafted filter banks. This approach was followed for speech recognition (Zeghidour et al., 2021). The only application to EEG relied on real-valued wavelets which provides incomplete representation, e.g. hindering the computation of phase-related features (Barmpas et al., 2023). Moreover, real-valued wavelets lack the desirable property of being invariant to small shifts after taking the modulus.

**Riemannian geometry** is a theoretical framework for analyzing symmetric positive-definite (SPD) matrices that forms the foundation of robust machine learning methods extensively applied to various prediction tasks involving EEG signals. These methods mostly focus on the processing of sample covariance matrices, which up to some regularization are SPD (Barachant et al., 2010, 2013; Congedo et al., 2017; Sabbagh et al., 2019). Such approaches present two major assets. First, they utilize an affine invariant metric which is relevant for EEG as the data-generating process can be modeled as a linear mixing of sources (Mosher et al., 1999; Nunez and Srinivasan, 2006; Sabbagh et al., 2020). Second, they project SPD matrices to a Euclidean tangent space, therefore enabling the use of classical statistical learning tools (Arsigny et al., 2007). Importantly, the tangent space mapping involves matrix logarithms which match well the dominance of lognormal distributions in brain structure and function (Buzsáki and Mizuseki, 2014). In practice, the log-Euclidean metric is often preferred to the affine invariant due to its superior computational efficiency while preserving theoretically desirable properties and yielding similar experimental results (Arsigny et al., 2006). The tangent space vector of a SPD matrix **X** is obtained by applying the logarithm map 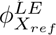 at the reference point **X***_ref_*,

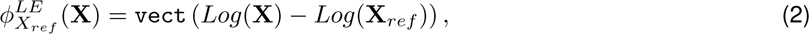

where *Log* is the matrix logarithm and vect is the vectorization operator that stacks the upper triangular elements of a matrix and multiplies the off-diagonal terms by 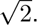.

Geometric operations for processing SPD matrices have recently been integrated into deep neural networks with mostly four operations (Brooks et al., 2019; Huang and Gool, 2017): The bilinear mapping, the rectified eigenvalue, the logarithm of eigenvalues and the batch normalization. Nevertheless, when applied to EEG signals, these operations present two main limitations that we aim to address in this work. First, when processing sample covariance matrices in the context of EEG, a well documented hurdle is the rank-deficiency induced by EEG preprocessing (Absil et al., 2008; Sabbagh et al., 2019). This issue is often ignored or ad-hoc mitigation approaches are used such as selecting a limited number of channels or adding a scalar to diagonal terms (Carrara et al., 2024; Wilson et al., 2022). Second, the projection to tangent space requires a reference point (**X***_ref_* in Equation 2), for which the mean across subjects is an intuitive choice as it reveals high-amplitude spatially structured noise related to volume conduction and anatomy-driven signal mixing. Intuitively, this allows computing power ratios relative to background activity (Nikulin et al., 2011; Sabbagh et al., 2020).

One can note that in DL models, logarithm maps use the identity matrix as reference rather than the average. When setting the reference to identity, the log map operation simplifies to the matrix logarithm, which reduces the computational and engineering complexity. However, this may not be optimal for EEG applications as the spatial patterns captured by between-electrode covariance matrices are governed by structured noise reflecting volume conduction (Engemann and Gramfort, 2015). Incorporating biophysically meaningful reference points such as the average covariance therefore enables explicit signal denoising. Learning the reference point was, therefore, an important step for matching the mathematical operations and the performance of non-deep Riemannian models (Bomatter et al., 2023; Sabbagh et al., 2020).

Taken together, the relevance and successful application of wavelets for spectral analysis and Riemannian geometry for prediction task suggest integrating these elements for neuroscience-informed DL architectures.

## 3 Methods

### 3.1 Model

In this work, we developed a novel architecture for the end-to-end processing of raw EEG data combining the extraction of time-frequency features with learned filter banks of Gabor wavelets, geometric transformations and fully connected layers (Figure 1). In the equations presented in following paragraphs, **X** (with additional upper or lower scripts and dimension specification) is used to describe the input of different layers but does not refer to EEG signals.

**Figure 1:**
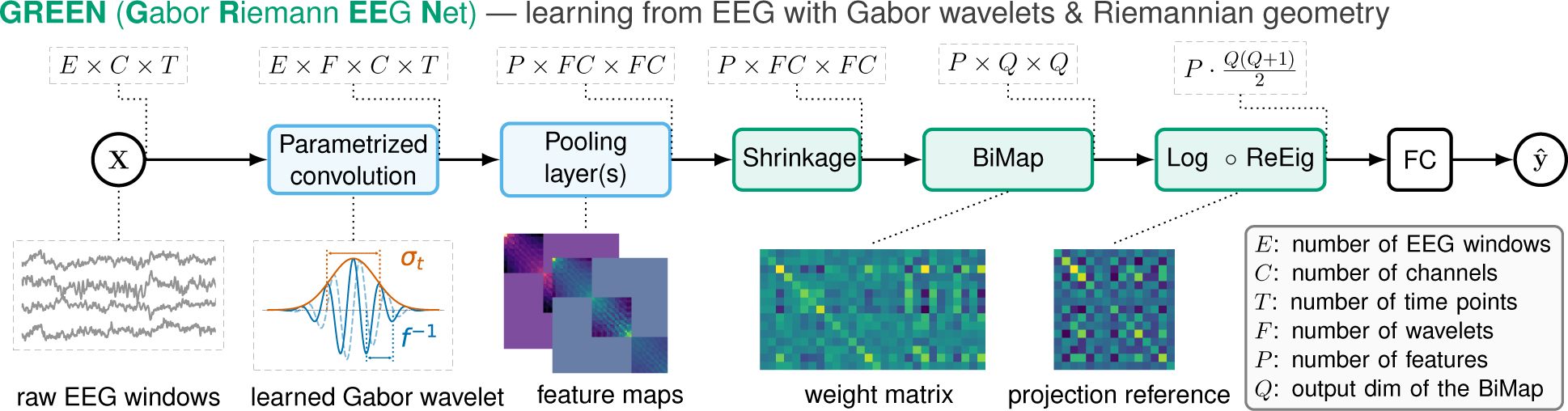
Composable *GREEN* building blocks and model architecture. The input consists of randomly drawn windows from EEG recordings (*E* windows, each with *T* time points). Blocks colored in blue operate on inputs that have a temporal dimension, whereas blocks colored in green operate on matrices. In the first convolution block, the kernel is a parametrized complex-valued Gabor wavelet. The parameters of this convolution layer are the standard deviation *σ_t_* of the Gaussian window and the central frequency *f* of the complex sinusoid. The pooling layer comprises the computation of the covariance matrix from the wavelet-transformed signal. Using equal wavelet lengths ensures the consistency of wavelet-transformed signals and allows us to evaluate cross-frequency interactions between channels. The pooling layer can accommodate other features derived from wavelet-transformed signals e.g. pairwise phase-locking value (the number of features being denoted *P*). The following layer performs a shrinkage operation on the symmetric positive semi-definite feature matrices, leading to down-weighing off-diagonal adapting a Ledoit-Wolf scheme (Ledoit and Wolf, 2004) where the shrinkage parameter is trainable. The subsequent bi-linear mapping (BiMap) and rectified eigenvalue (ReEig) layers have been introduced by (Huang and Gool, 2017) and provide computations on the Riemannian manifold including spatial filtering, dimensionality reduction and eigenvalues non-linearity. We modified the original implementation of the LogEig layer. Instead of the identity, a log-Euclidean running mean is used as reference point for the projection. This provides a meaningful reference point accounting for the dominance of spatial patterns induced by electromagnetic field spread and volume conduction. Finally, the tangent space vectors are fed to fully connected (FC) layers that can be designed and adapted for specific modeling goals.

#### Parametrized convolution

The first block of this architecture is the parametrized convolution layer. The kernel consists of complex-valued Gabor wavelets (Equation 1) in which the frequency of the carrier (*f*) and the standard deviation of the Gaussian window (*σ_t_*) are learned parameters. These time-frequency atoms can be interpreted as bandpass filters. As shown in Figure 1, for each sample, multiple time windows (*E*) are processed by this layer. Preliminary experiments showed that using *T* =10-second time windows led to good results in the explored range. For the initialization of the parameters *f* and *σ_t_*, we used the Morlet parametrization described above unless explicitly stated that a random initialization was used.

#### Pooling layer

The wavelet transformed signal is then passed through a pooling layer in order to compute feature maps. In the basic configuration, the pooling layer includes the sample covariance matrix computed from the wavelet-transformed signals. Concretely, given two wavelet-transformed signals 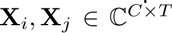 referring to wavelets *i^th^*and *j^th^* of the filter bank, the covariance matrix is

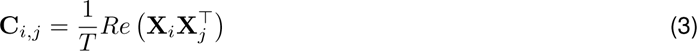

This leads to a matrix in which diagonal blocks contain the interactions between channels at a given frequency **C***_i,i_*, whereas off-diagonal blocks contain cross-frequency interactions between channels, **C***_i,j_*. Of note, in the present setting, the cross-frequency covariances represented by off-diagonal blocks can be expected to vanish. While we did not systematically explore this architecture component in the present work, this opens various opportunities for designing pooling layers for cross-frequency coupling (Canolty and Knight, 2010; Jensen and Colgin, 2007).

The complex wavelet-transformed signal allows for the computation of other features such as the pairwise phase-locking value (Lachaux et al., 1999). The pairwise PLV matrix **P** contains the elements

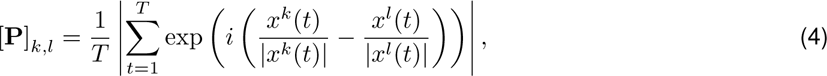

where (*k, l*) is a pair of sensors and *x^k^*(*t*) is the instantaneous value of the wavelet transformed signal. Dividing by the respective modulus provides a measure of the instantaneous phase difference, which is then averaged over time to compute the PLV. Besides these two examples of covariance and phase-locking that we study in the section 4 other features such as the power envelope correlation matrix (Hipp et al., 2011) or coherence (Varela et al., 2001) and various measures of cross-frequency relationships (e.g. phase-phase, phase-amplitude, amplitude-amplitude) could be used. Furthermore, the framework is applicable to the analysis of event-related brain activities if the EEG datasets subjected to the analysis are selected in an event-related manner.

#### Shrinkage layer

In the finite sample regime, sample covariance matrices are known to overestimate the range of eigenvalues which sometimes lead to numerical instabilities (Chen et al., 2010; Ledoit and Wolf, 2004). To mitigate this issue, we built on top of prior work on covariance shrinkage operators which are known to outperform ad-hoc approaches, especially in the EEG context (Engemann and Gramfort, 2015). Here, we implemented a shrinkage layer, which given a matrix 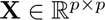 performs the operation,

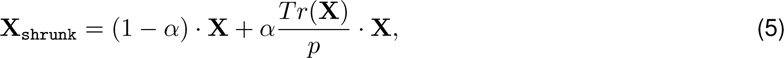

where the strength of the shrinkage *α* is a learned parameter. This operator presents the desirable property of preserving the trace of the matrix. For covariance matrices, this trace is the total power which is a critical feature of EEG signals often used as such in machine learning pipelines.

#### BiMap layer

In the context of EEG data, covariance matrices can be rank-deficient due to pre-processing and artifact removal (Bomatter et al., 2023) and therefore do not belong to the SPD manifold. To mitigate this issue, (Sabbagh et al., 2019) formally proved that projecting such matrices to a common subspace enables regression modeling on SPD manifolds with statistical guarantees despite rank-deficient covariance inputs. To incorporate these insights, the subspace projection can be implemented using a BiMap layer which given an input matrix 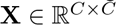 performs the operation,

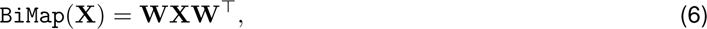

where the weight matrix 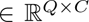 is constrained to be semi-orthogonal (Huang and Gool, 2017), using a manifold optimization algorithm (Lezcano-Casado, 2019), for more details, see subsection A.2. This step is consistent with classical spatial filtering (Dähne et al., 2014; de Cheveigné and Parra, 2014; Koles et al., 1990).

Empirical exploration of this approach confirmed that the eigenvalues of the projected feature maps were full-rank. Importantly, we found that this approach was also more robust to rank-deficiencies induced by preprocessing steps such as independent component analysis (Hyvärinen and Oja, 2000), which is practical as that approach allows to avoid the search for the smallest common subspace (Bomatter et al., 2023; Sabbagh et al., 2019). Furthermore, we observed that not constraining weight matrices to be semi-orthogonal still led to mappings that ensured the strict positivity of the resulting features maps, hence, fully supporting Riemannian computation. Therefore, to make computation more efficient,unless specified, we did not use a constrained optimization algorithm for the weights of the BiMap layer.

#### Combined ReEig and LogMap layer

The rectified eigenvalues (ReEig) layer creates a non-linearity transforming eigenvalues. Besides, this operation forces the eigenvalues to be greater than a given threshold, therefore ensuring numerical stability during training. Finally, the logarithm map (LogMap, see Equation 2), initially introduced as LogEig, uses the log-Euclidean metric which empirically provides similar results as the affine invariant metric at a cheaper computation cost (Arsigny et al., 2006). However, we differentiate our approach from others that use the identity matrix as reference. Instead, we use a running log-Euclidean mean **X***_ref_* which preserves the consistency with the Riemannian distances (Chevallier et al., 2021). The operation is provided in Equation 2. The update rule of the running mean at step *k* is given by,

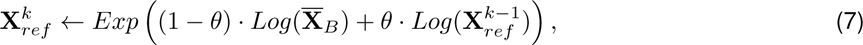

where *θ* is the momentum and 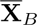 is the log-Euclidean mean over the current batch. *Log* and *Exp* respectively correspond to the matrix logarithm and exponential. Other works also explored Riemannian batch normalization but we did not observe any benefit of such a layer in our experimental setting in our past work (Brooks et al., 2019). To avoid redundant computation, we merged the last ReEig and LogMap computations by rectifying the eigenvalues and computing their logarithm in a single diagonalization step.

#### Fully connected layer

Finally, a classical multi-layer perceptron head can be added along other deep learning layers. Initial hyperparameter exploration led to good results with two fully connected layers (64 and 32 hidden units) with batch normalization and dropout (*p* = 0.333) and a GELU (Gaussian Error Linear Units) non-linearity (Hendrycks and Gimpel, 2016). For regression and classification tasks we respectively used the mean-square-error and the cross-entropy loss functions.

### 3.2 Datasets and preprocessing

We used three public datasets with institutionally controlled access. Our focus was on subject-level prediction using resting-state EEG which is most commonly used in clinical settings and biomarker applications. Single biomedical outcomes of human participants are predicted from the corresponding EEG recordings and the training and testing sets form a population of participants. This setup differs from the popular event-level prediction from event-related activity which is used in BCI where outcomes are defined for specific events e.g. stimuli or behavior. We therefore considered larger biomedical EEG datasets (more than 1000 participants) with published benchmarks for deep and non-deep ML models across diverse regression and classification problems. Finally, the selection of datasets covers a heterogeneous population, with healthy participants and patients suffering from psychiatric or neurodegenerative disorders, hence, allowing us to gauge the versatility of our approach.

#### TUAB

The Temple University Hospital Abnormal (TUAB) dataset contains resting state EEG recordings from 2993 patients with normal and abnormal labels provided by medical experts (Harati et al., 2014; Obeid and Picone, 2016). The recordings were acquired using variable numbers of electrodes (24 to 36). We used only the 21 channels that were common to all subjects and conformed to the 10-20 configuration. For the pathology classification task, we used all subjects and for the age prediction task, we considered a subset of 1278 healthy subjects to obtain results comparable with those published in previous works (Engemann et al., 2022; Gemein et al., 2023). Similarly to our previous work (Bomatter et al., 2023), we cropped recordings to a common duration of 15 minutes.

#### TDBRAIN

The Two Decades Brainclinics Research Archive for Insights in Neurophysiology (TDBRAIN) database contains 1274 EEG recordings of patients with psychiatric conditions (van Dijk et al., 2022). All recordings have been acquired using 26 electrodes positioned according to the 10-10 international system. Annotations indicated segments in which the subjects had their eyes open or closed are also provided. The dataset also contains demographic information including the sex of the patient (620 female, 654 male).

#### CAU

The Chung-Ang University (CAU) Hospital dataset contains recordings from 1155 patients along with clinical diagnoses (Kim et al., 2023). Subjects were grouped into three categories: normal, mild cognitive impairment and dementia. The recordings were acquired using 19 electrodes positioned according to the international 10-20 system. It should be noted that this dataset includes resting state but also EEG recorded during photic stimulation exposure periods. The average and standard deviation of the recording duration are respectively 13.34 and 2.83 minutes.

#### Common processing steps

For all the datasets the preprocessing included average referencing and band-pass filtering between 1-100*Hz*. High amplitude artifacts were removed using the autoreject algorithm (Jas et al., 2017). TUAB and TDBRAIN were resampled to 125*Hz* and CAU to 140*Hz* (half of the sampling frequency) to reduce the memory footprint. For the eyes-open versus eyes-closed classification task, we additionally included an ICA step to remove ocular artifacts that would otherwise be trivially predictive. We adopted the ICA pipeline as described in our work (Bomatter et al., 2023) based on the fast PICARD implementation (Ablin et al., 2018) of the FastICA algorithm (Hyvärinen and Oja, 2000) and the ICLABEL method (Li et al., 2022; Pion-Tonachini et al., 2019) for detection of artifact components. For all datasets and tasks, each sample consisted in 10 windows (*E* = 10) of 10-second (*T* = 10*s*) randomly drawn from the recordings.

### 3.3 Baseline model

In previous publications, methods based on Riemannian geometry have been reported to perform similarly to a wide range of deep learning architectures on the pathology classification task on TUAB (Gemein et al., 2020), and provided comparable results to a DL model on the age and sex prediction tasks on TUAB and TDBRAIN (Bomatter et al., 2023; Engemann et al., 2022; Sabbagh et al., 2020). Here we focused on the model presented by Bomatter and colleagues (2023) which we call baseline throughout this work. The *GREEN* architecture was directly inspired by this baseline, capitalizing on wavelets and incorporating equivalent operations in its shrinkage, BiMap and LogMap layers. Choosing this baseline allowed us to understand which precise additional computational steps and non-linearities can add improvements.

The baseline used features extracted from wavelet-transformed signals using a predefined Morlet filter bank (49 wavelets with central frequencies logarithmically spaced between 1 and 64 Hz). Sample covariance matrices were then estimated and projected to the SPD manifold using Principal Component Analysis (PCA) to account for their rank-deficiency (Sabbagh et al., 2019). Finally, a projection to tangent space (Equation 2) followed by a linear model with ridge regularization were applied (Hoerl and Kennard, 1970). As in the original work, we used the scikit-learn (Pedregosa et al., 2011) implementation RidgeCV, which uses generalized cross validation (Golub and von Matt, 1997) for hyper-parameter selection of the penalization strength. We tested 100 values between 10*^−^*^5^ and 10^5^ on a logarithmic grid.

To compare results obtained using other deep learning architecture, one the CAU dataset, we reproduced the exact train test-splits and reported the previously published benchmarks (Kim et al., 2023).

### 3.4 Statistical inference for model comparisons

Except for the pathology classification on CAU where the test set from the original publication was used for comparison (Figure 2b), we used a Monte-Carlo cross-validation scheme to compare the performance of the different models. The experimental protocol consist of 100 random splits with a test set containing 20% of the data as described by (Bouckaert and Frank, 2004). To make up for the underestimation of the variance in cross-validation settings, we followed Nadeau and Bengio’s corrected resampled t-test

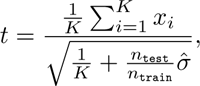

where *K* is the number of splits, *n*_train_*, n*_test_ are respectively the train and test sizes, *x_i_* is the performance difference between the two models and 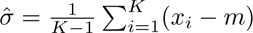 is the naive estimate of the variance, *m* being the mean performance difference.

**Figure 2:**
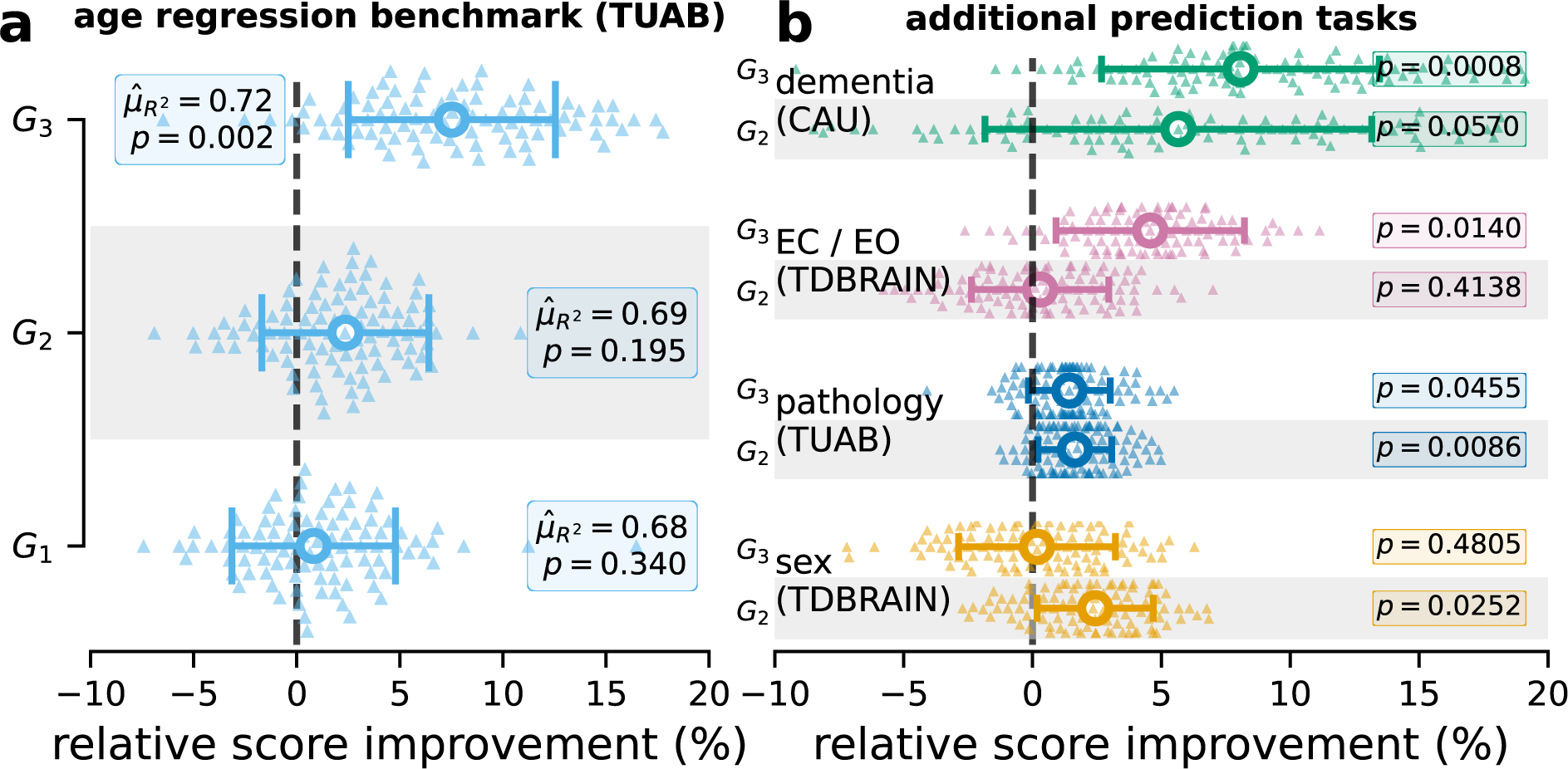
The impact of GREEN architecture components varied across prediction tasks. **(a)** cross-validated performance (100 Monte-Carlo splits, 20% test set size) for age regression on TUAB data (healthy subjects) for models of increasing complexity (*G*_1_ 3, cf. Table 1). Boxplots show distributions of percent changes of *R*^2^ between each model versus the baseline. Annotations show the absolute score of the model (*G*_1_ 3) and hypothesis tests (null-hypothesis being no improvement over baseline). *G*_1_ and *G*_2_ used the same filter bank as the baseline. *G*_1_ mirrors the non-deep baseline but uses a stochastic optimization procedure. *G*_2_ added a hidden layer enabling non-linear functions beyond the logarithm implied by the tangent-space embedding. In addition to the hidden layer, *G*_3_ also used trainable wavelets and takes raw EEG as input. *G*_3_ showed markedly higher performance than *G*_1_, suggesting that fine-tuning of frequency filtering and non-linearities were beneficial for this task. **(b)** same experimental protocol as in (a) for four additional tasks (from top to bottom): three-way classification of diagnosis (healthy, mild cognitive impairment (MCI), or dementia), binary classification of recordings with eyes open versus eyes closed, of pathological versus normal recordings and of the patient’s sex. For each task the dataset used is specified in parentheses. The advantage of the architectures *G*_3_ depended on the task and was most pronounced for dementia diagnosis and classification of eyes open (EO) versus eyes closed (EC).

**Table 1:** Architecture details for the different models studied.

| Input<br>Component | raw |  | feature map |  | vector |  | Adam |
| --- | --- | --- | --- | --- | --- | --- | --- |
|  | filter bank | pairwise PLV | BiMap | LogMap | linear | hidden layer |  |
| **Model** |  |  |  |  |  |  |  |
| baseline | X | X | X | ✓ | ✓ | X | X |
| $G_1$ | X | X | ✓ | ✓ | ✓ | X | ✓ |
| $G_1P$ | X | ✓ | ✓ | ✓ | ✓ | X | ✓ |
| $G_2$ | X | ✓ | X | ✓ | ✓ | ✓ | ✓ |
| $G_3$ | ✓ | ✓ | ✓ | ✓ | ✓ | ✓ | ✓ |
| $G_3P$ | ✓ | ✓ | ✓ | ✓ | ✓ | ✓ | ✓ |

## 4 Results

### The impact of architecture components depends on the prediction task

We first explored different architecture components and evaluated their performance on five tasks and three datasets with available benchmarks (Figure 2). The five benchmarks include age regression on the TUAB dataset, binary classification of pathological versus normal recordings using the TUAB dataset, binary sex classification on TDBRAIN dataset, binary classification of recordings with eyes open versus eyes closed from the TDBRAIN dataset, and, three-way classification of diagnosis (healthy, mild cognitive impairment (MCI), or dementia) using the CAU dataset. By design, the modularity of the *GREEN* architecture supported this objective by allowing us to remove or freeze the parameters in order to reproduce the operations performed by baseline methods and assess the benefits of additional complexity (cf. Figure 1).

We first revisited age prediction on the TUAB (Figure 2) which has been intensely studied with Riemannian filter bank regression in prior work (Bomatter et al., 2023; Engemann et al., 2022; Sabbagh et al., 2020). Results are shown for three different models, *G*_1_ reproduces the operations of the baseline, including a pooling layer that computes only the covariance (*P* = 1), and a BiMap layer but uses a stochastic optimization method. The input dimension of the BiMap depended on the number of channels *C* (see subsection 3.2) and the output dimension was set to *Q* = *C* − 1. *G*_2_ has an additional hidden layer (32 units). *G*_3_ uses a learnable filter bank of *F* = 10 wavelets and two BiMaps of respective output dimensions *Q* = 64 and *Q* = 32. For more details refer to Table 1. Our results showed that the addition of a hidden layer improved performance by a few percent (not statistically significant). This comparison is meaningful to empirically test the logarithm hypothesis motivated by insights into log-dynamic scaling of brain structure and function (Buzsáki et al., 2013; Sabbagh et al., 2020). For the case of age-prediction, results suggests that the logarithm is probably a sufficient modeling assumption to solve the age prediction task. Importantly, the greater improvement was observed for model *G*_3_, which introduced filter banks of trainable wavelets, suggesting that task-relevant information can be better captured with learned filters rather than a fixed grid.

We extended the comparison between the baseline, *G*_2_ and *G*_3_ architecture to the four remaining classification tasks (Figure 3 b). The results showed that the relative improvement over the baseline varied depending on the task, confirming the intuition that the spread of information along the frequency spectrum may be task-dependent. Improvements with *G*_2_ were significant (*α* = 0.05) for sex prediction and pathology decoding. Improvements with *G*_3_ were significant (*α* = 0.05) for eyes-closed versus eyes-open classification, pathology decoding and dementia classification. For a detailed report of performance scores and test statistics, see Table S1.

**Figure 3:**
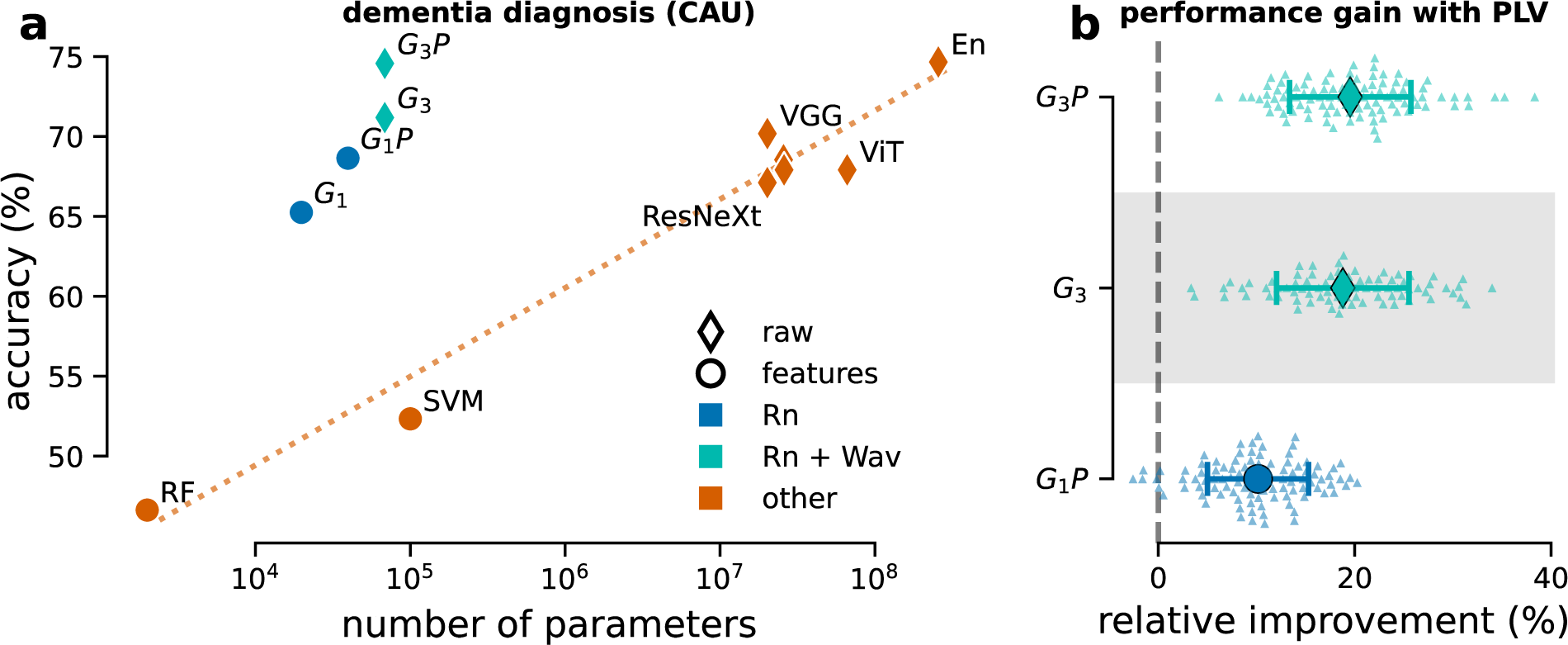
GREEN performance-complexity trade-off across components and extensions. **(a)** model performance versus complexity measured with the number of parameters for the three-way classification of dementia diagnosis on the CAU dataset. The accuracy is measured on the benchmark from Kim et al. 2023 using the same single train-test split. Results depicted in orange were replotted from the original publication. Results in blue show proposed models based on Riemannian geometry (Rn). Cyan represents the combination of Riemannian geometry and learned wavelets (Rn + Wav). Next to covariances, *G*_1_*P* and *G*_3_*P* also included pairwise phase locking matrices to capture the phase information that might arise from the photic stimulation periods present in the EEG recordings from this dataset. It can be seen that the Riemannian models with learnable wavelets and explicit phase-locking features achieved high performance with orders of magnitude fewer parameters. **(b)** performance gain over the baseline model *G*_1_ obtained by including PLV matrices in the pooling layer (*G*_1_*P*), learnable wavelets (*G*_3_) and the combination of both (*G*_3_*P*). The same experimental protocol as in Figure 2 (100 shuffled splits, 20% test set size) was used. Including PLV matrices as a measure of phase synchrony in the pooling layer improved performance significantly for *G*_1_*P* over *G*_1_ and slightly for *G*_3_*P* over *G*_3_.

The benefits of learnable wavelets was most pronounced for dementia and eyes-closed versus eyes-open classification as indicated by cross-validation uncertainty and average performance differences. Furthermore, a predefined filter bank can lead to similar (pathology decoding) or even better (sex prediction) performance than a learned filter bank and, likewise, the more complex FC layers can outperform the logarithm nonlinearity (sex prediction). These results emphasizing the importance of a modular approach to build models that can be adapted to the task at hand.

### *GREEN* shows a favorable performance-complexity trade-off while remaining extensible

The CAU dataset (Kim et al., 2023) offered us the opportunity to position the proposed models on a benchmark for diagnosis prediction in dementia against recent large DL architectures (Figure 3 b). Replotting the results from Kim et al. 2023 revealed a log-linear scaling of performance with model size, in number of parameters (Pearson correlation coefficient of 0.98). In contrast, proposed models based on Riemannian geometry (Rn, model *G*_1_*, G*_1_*P*) seemed to break this trend and presented a favorable performance-complexity trade-off. We observed that increasing the complexity of the model by allowing the learning of the parameters of ten wavelets (Rn + Wav, models *G*_3_ & *G*_3_*P*), the performance was largely improved, outperforming the best single model presented in the original paper and matching the performance of the ensemble model (bag of all the other models presented in the paper) (Figure 3 a). These performance gains were achieved with up to three orders of magnitude fewer parameters than DL other DL models.

Availability of prior information that the recordings contained segments of photic stimulation, which has the capacity to alter neural phase-synchrony in stimulation (Infantosi and de Sá, 2006; Lakatos et al., 2019; Schroeder and Lakatos, 2009), motivated us to add a widely used measure of phase synchrony i.e. pairwise PLV (Lachaux et al., 1999) matrices in the pooling layer (see Equation 4). The lean design of the proposed *GREEN* architectures allowed us to investigate the hypothesis regarding the relevance of phase-locking features more rigorously by running model comparisons with full-blown Monte-Carlo cross-validation scheme with 100 repetitions (unfortunately not available for the original CAU benchmark). When concatenating the covariance matrices with PLV matrices the performance was improved significantly for *G*_1_*P* over *G*_1_ and slightly for *G*_3_*P* over *G*_3_ (Figure 3 b), suggesting that phase-locking measures were informative but not indispensable for the more complex architecture with trainable wavelets.

These experiments evidence two key features of the GREEN architecture. First it improves the performance-complexity trade-off by integrating prior knowledge in the architecture, through parameterized wavelets and Riemannian computations which alleviates the size of models without reducing the prediction accuracy. This allows to outperform large DL models with orders of magnitude fewer parameters. Second, the modularity of the framework facilitates the integration of classical neuroscience measures of phase synchrony. This facilitates testing of specific scientific hypotheses regarding the predictive importance of e.g. phase-synchrony features in the context of EEG recordings containing photic stimulation periods.

### The optimal numbers and frequencies of wavelets depends on the prediction task

Our results revealed how the proposed architecture can lead to performance benefits at a low computation cost. In this context, the size of the filter bank is an intuitive hyperparameter, measuring the spread of the information along the frequency spectrum. This raises the question to which extent task-relevant information is concentrated versus distributed along the frequency spectrum. The filter bank that presents the optimal trade-off between size and performance could thus be seen as a measure of the model complexity needed to solve a task.

We investigated the minimal filter bank size needed to reach the performance of the most complex model. For each task, we progressively increased the filter bank size using *F* = 1*,…,* 15 wavelets and tracked the performance on the same single testing fold, error bars are measured for five random initializations (Figure 3 a). For each task, the curve was plotted until the model achieved a performance less than one standard deviation away from the model with the largest filter bank (*F* = 15), which consistently yielded the best results. It can be seen that the number of wavelets needed to reach this best performance was highly task-dependent, providing insights into the model complexity requirements.

The results for the dementia diagnosis and the eyes state tasks revealed that the minimal filter bank size was only two or three wavelets. On the other hand, for age- and sex-prediction, more wavelets were required, which, in the case of sex prediction did not transform into improved performance over the baseline. This suggests that the predictive spectral signatures on certain prediction tasks were more localized and less diffuse in the frequency spectrum. In these cases, the *G*_3_ learned to extract sparse representations of the complex EEG signals that concentrated the information needed to solve prediction tasks.

We then inspected the center frequencies of the wavelets learned by the model with a sparse filter bank for each of the five tasks studied (Figure 4b). Such frequencies could be interpreted as the most predictive and least redundant spectral features of the signal. Across tasks, the low frequency region of the spectrum was often selected alongside the alpha band (8-12*Hz*), which is a strong signature of wakeful human EEG and has been intensely studied due to its favorable signal-to-noise ratio.

**Figure 4:**
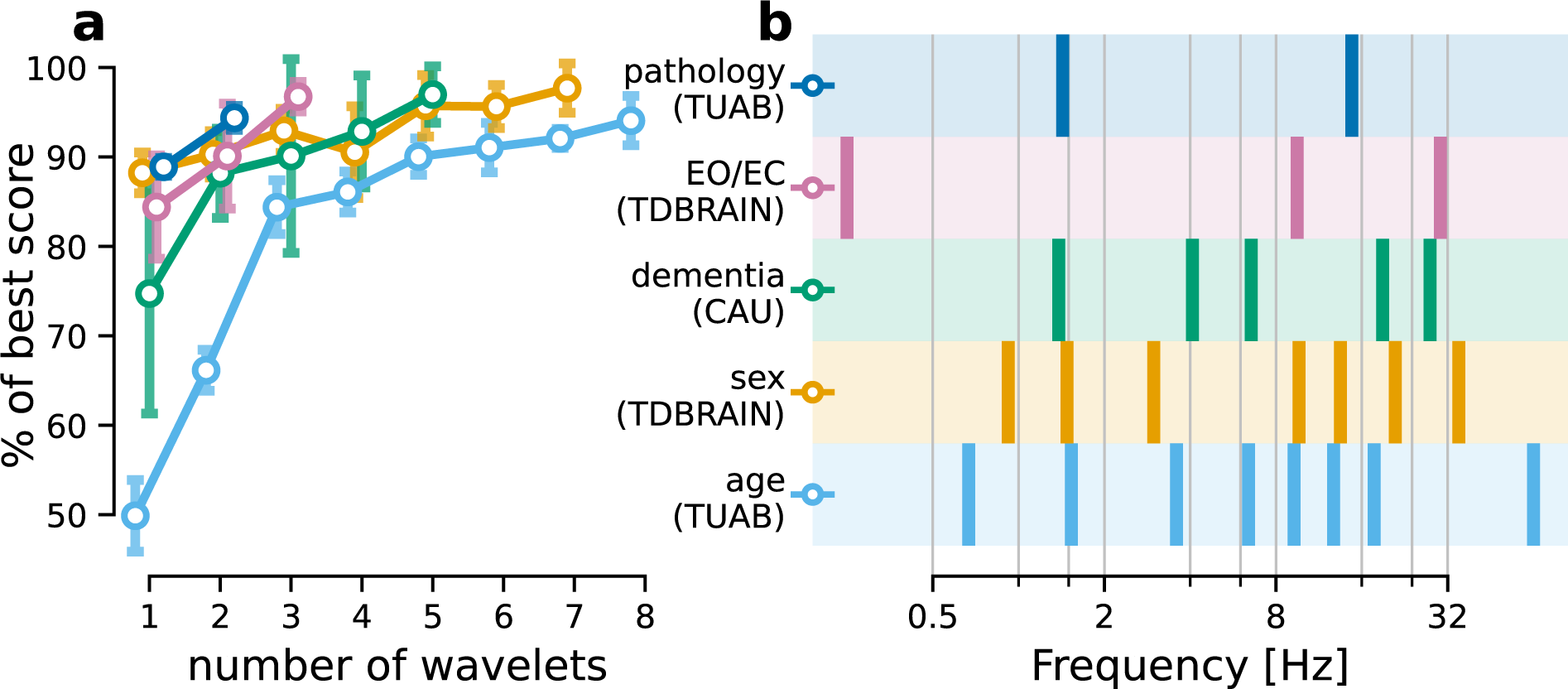
Optimal filter bank composition depended on the prediction task. **(a)** model performance as a function of model complexity in terms of numbers of wavelets. Performance for all classification tasks and age regression (Figure 2) is presented as a percentage of the asymptotic value (best model) measured using a filter bank of 15 wavelets. Curves were plotted until the mean performance is less than one standard deviation away from the final asymptotic performance. Uncertainty estimates were obtained by repeating the analysis five times using random initializations. It can be seen that the number of wavelets required substantially varied across tasks and can be surprisingly low with two to three wavelets only i.e. for pathology and EO/EC classification. By comparison, the baseline (cf Table 1) used a filter bank of 49 wavelets. (**b**) frequencies of Gabor wavelets in the filter banks learned by the model for the different tasks. The number of wavelets was selected based on the learning curve presented in (**a**). Precise frequencies varied across tasks, however, low frequencies between 1*Hz* and 2*Hz* and in the alpha range (8-12*Hz*) were common choices.

**Figure 5:**
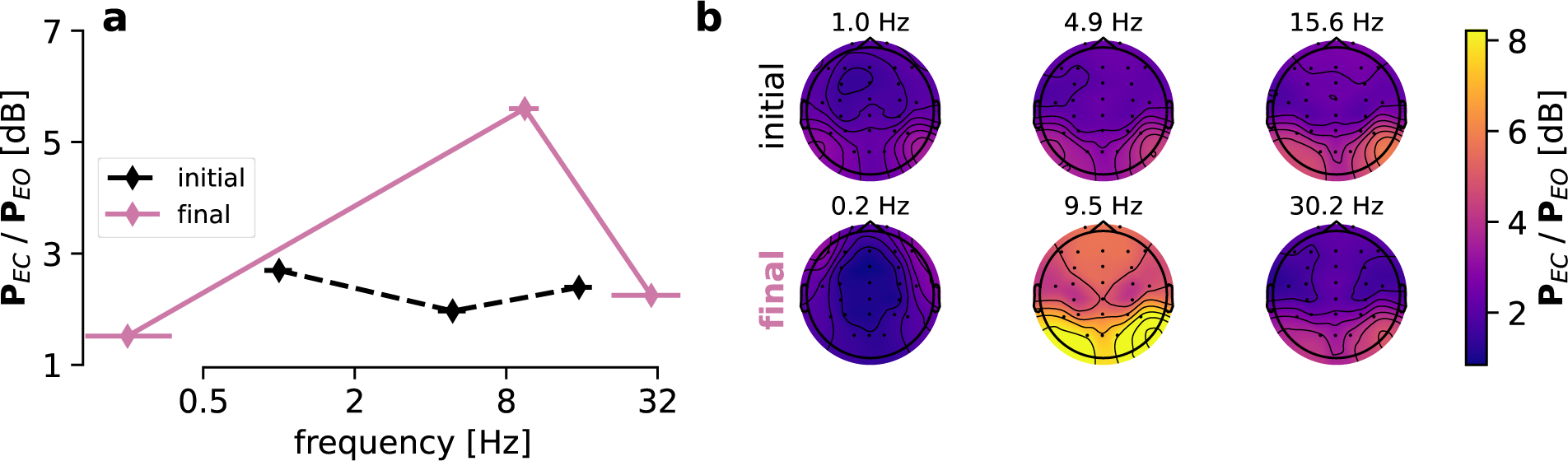
Task-specific learned representations provide actionable EEG quantities. (**a**) averaged EEG power ratio (EC/EO) in dB derived from the wavelets transformed signal, using either the randomly initialized model (dashed line) versus trained (solid purple line). Horizontal error bars indicate the frequency domain standard deviation of the wavelet. The model focused on 8 − 12*Hz* (typical location of occipital alpha activity with eyes closed) as well as low and high frequencies as potential reference points. Markedly, after training, the frequency-domain standard deviation was minimal for the wavelet with the precise frequency near 9*Hz* which differs from commonly used parametrizations. (**b**) topographies for the three wavelets before (initial) and after (final) training. Powers obtained from the central wavelet showed an occipital topography characteristic for idling visual alpha rhythms. These power topographies were obtained from a forward pass (corresponding to the diagonal of the covariance matrix) and can be readily reused for secondary statistical analysis or visualization.

These results highlights a strength of the proposed architecture to uncover the informative frequency ranges via supervised learning. To the contrary, classical pipelines rely on pre-defined filter banks and require a far more fine grained coverage of the frequency spectrum to solve certain tasks, leading to larger filter banks. In sum, such profiling benchmarks can provide important leads for in-depth exploration of underlying predictive EEG signatures and task-specific markers.

### *GREEN* yields interpretable and actionable EEG quantities for biomarker exploration

The modular design of *GREEN* which is based on scientifically interpretable components encourages further in-depth exploration and extraction re-usable EEG quantities (Figure 4). As wavelets can be interpreted as bandpass filters it can be seen that the diagonal of the covariance matrix contains classical EEG-power topographies. To investigate the utility of GREEN for data exploration we investigated the eyes-closed versus eyes-open prediction task in more detail. Since Hans Berger’s discovery of the EEG, it has been known that closing the eyes predictably intensifies occipital alpha rhythms that have been linked across species to sensory idling functions and “partial disengagement from the environment” (Buzsáki, 2006). This well-known phenomenon is thus a good toy example to study the utility if the *GREEN* architecture to probe physiological understanding. It can be seen that *G*_3_ places one wavelet exactly in the expected 8-12*Hz* region, where the power difference is most visible (Figure 4). In addition, model selection showed that three wavelets were needed to achieve high performance. It is noteworthy that the frequency domain standard deviation is minimal for the wavelet located in the alpha band. This is a substantial deviation from the popular Morlet parametrization that log-linearly increases the spectral smoothing with frequencies and, therefore, suggests that a more precise localization in the frequency domain is beneficial for this task. As this task was performed across subjects, and alpha rhythms are known to show high individual variability, one plausible explanation could, therefore, be that *G*_3_ might have used the low- and high-frequency wavelets as reference points to compute power ratios isolating alpha rhythms from their broadband background activity. Finally, inspecting the topography of the central wavelet after model training revealed, a typical visual-occipital alpha pattern emerged (Figure 4 c). It is important to highlight that these visualizable EEG quantities were fully derived from the activations of the trained *G*_3_ network without further external computation.

These results illustrate how the *GREEN* architecture can be practically used for biomarker exploration by extracting EEG descriptors such as power values and topographies that are well-established for traditional statistical visualization and scientific hypothesis testing.

## 5 Discussion

The present work introduced a novel lightweight DL architecture termed *GREEN* (Gabor Riemann EEGNet) for predicting and learning from EEG in a unified framework. By integrating wavelets in a DL architecture our approach provides a flexible and interoperable tool for probing modelling assumptions, exploring EEG data and predicting with potential state-of-the-art performance. Furthermore, the relatively small number of parameters makes it easy to train this model with limited computation resources hence fostering reproducibility and future research works.

### A novel interoperable light-weight neural network for prediction of biomedical outcomes from EEG

We tested the *GREEN* model on five different prediction tasks and three datasets with EEGs from more than 5000 human participants. *GREEN* uses convolution with parameterized Gabor wavelets which presents a twofold benefit. First, they can easily be interpreted as bandpass filters and allow for the computation of features that have been extensively studied by the neuroscience community but may be overlooked by ML practitioners. Second, this parametrization is characterized by only two parameters, hence drastically limiting the total number of parameters in the architecture in comparison with other DL approaches. Furthermore, this architecture supports adequate geometric operations on the manifold of SPD matrices, therein providing support for a broad set of data descriptors including covariance, power-envelope correlation matrices and other bi-variate interaction measures used in the literature for estimating functional connectivity in cortical networks (Hipp et al., 2012; Sadaghiani et al., 2022). The set of pooling layers is made even larger by the possibility to measure cross-frequency interactions (Canolty and Knight, 2010; Jensen and Colgin, 2007). This was not possible using common implementations of the Morlet parametrization, in which wavelets have varying temporal span depending on the frequency (Morlet et al., 1982), leading to discrepancies in the dimensions of the wavelet-transformed signals. While such terms typically vanish in the case of covariance matrices the phase of the transformed could be removed by taking its modulus in order to compute the power-envelope correlation matrices. This new lightweight models shows state-of-the-art performances in established benchmarks such as age regression, sex prediction or pathology classification with orders magnitude less parameters than baseline DL architectures.

### A modular framework for testing modeling assumptions

Our work demonstrates how the modularity of deep learning can be used to express and investigate hypotheses about the underlying mechanisms relating EEG signals to biomedical outcomes. We designed the *GREEN* model to integrate established concepts from neuroscience and EEG e.g. neural oscillations, power and phase-synchrony with Riemannian geometry and deep learning methods through one single model. As the optimization engine is held constant across model comparisons, this approach provides high granularity for testing the impact of specific components.

Concretely, this has led us to the insight that certain tasks could be better solved by a few learned wavelets than a pre-defined grid of wavelets. Moreover, we found mixed evidence for the benefit of more complex non-linearities beyond the logarithm modeling assumption (Buzsáki and Mizuseki, 2014; Sabbagh et al., 2020) as the added value of the fully-connected layer was only visible for sex prediction and pathology decoding but not for the other prediction tasks. Finally, our approach facilitates the integration of additional feature computation, as illustrated by the addition of pairwise phase-locking value matrices which we found to convey complementary information over the covariance for dementia diagnosis. At the methodological level, our work is related to recent non-deep Riemannian ensembling approaches in the BCI context which combined different types of EEG connectivity features –cast into SPD matrices– (Corsi et al., 2022) or stacked generalization (Wolpert, 1992) of sub-models for specific M/EEG features in the context of age prediction (Engemann et al., 2020; Sabbagh et al., 2023). Another recent study proposed a Bayesian method extending the tradition of ICA for unsupervised learning of oscillatory components, resembling our work in providing inference on relevant frequencies (Das et al., 2023). However, we would argue that one distinctive feature of our proposed framework is coherent computations using the same wavelet bases guided by one single supervised optimization objective. Taken together, the proposed framework supports a bottom-up approach to building deep learning models in which trusted components are gradually combined and tested rather than applying entire DL models developed in other domains (vision, speech) to EEG.

### A potential workflow for EEG-biomarker exploration and practical considerations

The modular design and the choice of components mirroring concepts from the neuroscience literature has the potential to facilitate EEG-biomarker exploration in clinical applications. As one concrete toy example we demonstrated for the well-studied Berger effect (closing eyes inducing 8-12Hz oscillations) how frequency and EEG power can be directly extracted from a trained *GREEN* model via its forward pass. Although this example focused on a well-studied phenomenon, it allowed us to uncover surprising elements, for example the importance of low and high frequencies for detecting if eyes are closed or open. Interestingly, inspection of the frequency-standard deviation of the wavelets revealed a non-monotonic function with a more localized wavelet in the expected alpha-band range around 10*Hz* and broader frequency spread for wavelets located at low and high frequencies. It is noteworthy that this type of pattern reflects a unique capability of the proposed architecture to learn a flexible filter bank parametrization that is not bound to traditional rules such as increasing the frequency-standard deviation log-linearly with increasing frequencies as e.g. Morlet wavelets (Bomatter et al., 2023; Hipp et al., 2012; Morlet et al., 1982; Tallon-Baudry et al., 1996). Such deviations can be scientifically exploited as they convey insights into the data.

To continue within this example, we found the consistent choice (across five random initializations) of the 0.5*Hz* low-frequency wavelet to be surprising as the data was high-pass filtered at 1*Hz*. Additional inspection of the intermediate representations of an individual EEG recording Figure S1 revealed that, as expected given the filtering, the amplitude of the 0.5*Hz* wavelet-transformed signal was an order of magnitude lower. Surprisingly, we observed physiological oscillatory activity, pointing at information in low frequencies that was relevant for prediction and therefore recovered by the model from the attenuation of the high-pass filter.

These observations lead us to argue that representations learned by the *GREEN* architecture facilitate scientific exploration by readily providing established EEG descriptors such as power estimates that encourage visual and statistical comparisons. This practical level of interpretability is in particular important given the regulatory constraints on statistical analysis of biomarkers. These require that biomarkers can be reproducibly measured, which, e.g. in a large multi-centric clinical trial is operationally more likely to succeed with simpler procedures. Furthermore, biomarkers must stand in a mechanistic relationship to the disease biology and targeted physiological pathways, which has been repeatedly established for EEG power in specific examples (Frohlich et al., 2019a,b; Hipp et al., 2021; Janz et al., 2022). In this context, ML could assist EEG-biomarker development through at least two main routes, which we shall stylize for clarity: *Route A* uses learned representations to generate novel insights into the data and inform an explicit rule-based, operational biomarker recipe. *Route B* directly uses model predictions or learned representation as biomarker values. Importantly, the *GREEN* architecture provides high flexibility for pursuing any of these routes. However, what makes it different is that its representations have names in the neuroscience literature and that its lean design facilitates rapid exploration cycles of model training and subsequent visual or statistical exploration. We argue that these properties should largely improve the capacity of *GREEN* to support *route A* while its high performance as a prediction engine maintain access to *route B* without compromises.

This brings us to one more consideration. In our benchmarks, *GREEN* revealed its potential for high prediction capacity enabled by structural priors and constraints on computational complexity. It is fast to train and therefore supports scientific agility, not only translating into more exploration in less time and for lower electricity costs Thompson et al. (2024, 2020) but also removing obstacles for rigorous statistical inference practices based on resampling procedures. We see this as an important property for a scientific tool in empirical disciplines dealing with high-dimensional data (such as neuroscience) in which core discoveries are often driven by serendipitous exploration. Last but not least, a lean, efficient and scientifically interoperable modeling pipeline contributes to the democratization of computation (Hayashi et al., 2024; Renton et al., 2024).

Finally, the *GREEN* framework should be seen as a complementary tool rather than a drop-in replacement for previously proposed wavelet toolboxes (Bomatter et al., 2023). Indeed, learning filter banks in a supervised way can be a starting point for further analyses. A second step might consist in using a more fine grained grid in the identified frequency ranges or computing additional features from the wavelet-transformed signal. Another approach that was presented in this work is to use predefined filter banks as an initialization of the convolution layer and assess to which extent the model deviates during training.

### Theoretical considerations

We wish to point out that, beyond these practical applications, the proposed *GREEN* framework possesses favorable properties for future method development in mathematical modeling to study data-generating mechanisms. First of all, *GREEN* performs operations that are consistent with prediction models that benefit from desirable theoretical properties that were investigated formally in prior work (Sabbagh et al., 2019, 2020). This is eminently true for *G*_1_ which was designed to reproduce previously proposed pipelines. For this model, prior work (Sabbagh et al., 2019) showed that it enjoys the property of *statistical consistency* under the assumption that the outcome is generated by a linear combination of the log-power of stationary EEG sources in the presence of linear mixing between sources and electrodes. Notably, the theoretical analysis in Sabbagh et al. 2019 only treats the spatial dimension of the problem and assumes some arbitrary bandpass filter, hence, there is no loss of generality by the introduction of learnable wavelets. Furthermore, we would argue that the introduction of additional hidden layers in (*G*_2_ − *G*_3_) preserves this theoretical analysis. In fact, we found limited evidence that the hidden layers alone led to changes in the models’ performance (Figure 2).

This brings us to a final consideration. Despite added complexity and expressiveness, the SPD representations learned in pooling layers and BiMaps can be reused with non-deep approaches. For instance, as inputs for methods from the rich literature of kernel methods (Berlinet and Thomas-Agnan, 2011), which can accommodate manifold-valued data (Jayasumana et al., 2015), hence, enabling various applications in high-dimensional statistical learning (Sejdinovic et al., 2013; Wahba, 1981; Williams and Rasmussen, 1995). These properties might facilitate future work on developing uncertainty estimation and statistical inference within *GREEN* framework.

### Limitations & future work

*GREEN* unlocks performance with a lean architecture design enabled by the choice of strong structural priors such as sinusoidal complex wavelets and Riemannian geometry. This leads to characteristic inductive biases which allow sparse representations but may hit limitations. For instance, the ability of the wavelet convolutions to capture complex neural oscillations, e.g. documented cases of cross-frequency couplings, non-sinusoidal or asymmetric waveforms (Canolty and Knight, 2010; Cole and Voytek, 2017; Jackson et al., 2019), with a reasonably sized filter bank of wavelets requires experimental testing. We identified three potential extensions of the present work to tackle this challenging task. First, to capture and study non-sinusoidal waveforms, the filter bank could be constrained to contain groups of harmonic wavelets. Second, in an attempt to capture cross-frequency correlations, the pooling layers could also be extended to compute power-envelope covariances. Similarly, measures of phase-amplitude coupling could be considered. Another direction would be to iterate wavelet convolutions and modulus operator similarly to a scattering network (Bruna and Mallat, 2013). Such approaches are known to capture complex patterns allowing to discriminate between different signals that share a similar power spectrum. However, they typically use a fixed grid of wavelets and using a cascade of convolution would hinder the explainability of the model. Finally, this work studied Gabor wavelets due to their theoretically desirable properties, however, other waveforms from the rich wavelet literature (Mallat, 1999), that support such parametrization might as well be explored. In addition, while covariance matrices are useful for encapsulating the overall correlation structure between different channels over time windows, such representations are inherently limited in their ability to capture transient patterns. Non-stationary activities are however known to carry physiological information associated with certain conditions (Jing et al., 2020; Vidaurre et al., 2016). Mitigation strategies include working with distributions, instead of single covariance matrices, by using methods from the optimal transport framework (Bonet et al., 2023) or using sequences of inputs along with recurrent neural networks (Bashivan et al., 2015) or attention mechanisms (Yang et al., 2024).

This architecture enabled practical interpretability by allowing to carefully gauge the impact of network components on prediction performance. However, this is fundamentally different from statistical inference or variable importance estimation (Barber and Candès, 2015; Louppe et al., 2013; Nguyen et al., 2022). Recent extensions of permutation-based importance hold promise to provide statistical error control and effective variable-importance detection in the context of deep learning models and high-dimensional correlated inputs (Chamma et al., 2024a,b; Mi et al., 2021). The successful application of high-dimensional inference procedures will be an important opportunity for future work with the potential to add statistical rigor to the practical inference provided by the structural design choices of *GREEN*.

Finally, our proposed framework is yet to be explored in the context of interventional studies to address specific medical hypotheses or questions. Finding frequency bands and spatial patterns that are predictive of various endpoints, such as treatment response, disease progression or adverse events, remains a high-priority task for future applied work to inform biomarker development. In this context, self-supervised learning (Banville et al., 2021; Noroozi and Favaro, 2016) could be a promising extension of *GREEN* to tackle the unsolved challenge of cross-dataset transfer, for instance across centers. Even though EEG datasets from the general population keep increasing, for instance when considering advances brought by innovative multi-site consortia (Dzianok and Kublik, 2024; Haraldsen et al., 2024; Prado et al., 2023) in the dementia space, the number of patients included in clinical trials remains limited. Self-supervised approaches could, therefore, be an effective strategy for learning clinically robust representations that can be fine-tuned to solve data-set specific tasks and, at the same time, cope with covariate shifts (Dockès et al., 2021; Mellot et al., 2023). Concretely, the *GREEN* architecture could be used as the encoder of a more complex self-supervised learning architectures (Banville et al., 2021; Guetschel et al., 2024). As final limitation for clinical application, we have not studied the capacity of our model to adjust to high-density EEGs with 128 or more channels. The questions of how the model would predict from such data without modification and what would be the optimal size of the filter banks remain interesting open questions.

## 6 Conclusion

The new *GREEN* architecture was designed to facilitate the work of researchers working on biomarker technologies and clinical applications with EEG. Our results suggest that this effort to incorporate neuroscientific domain knowledge into the architecture design has the potential to offer new angles for approaching clinical problems. At the same time, this work is an interdisciplinary attempt to bridge biostatistical thinking in the context of clinical studies with machine learning research. The design-philosophy of *GREEN* reveal its own vision of DL for EEG in the era of large-scale models. Improving the effectiveness of well-known EEG signatures such as EEG power and phase-synchrony through strategic architecture design rather than perpetually building larger models following the “bigger-is-better” paradigm. To support this effort, we share the source code of *GREEN* ^1^.

## Author Contributions

In alphabetical order.

**Conceptualization**: D.E., J.F.H., J.P.

**Data curation**: D.E., J.P

**Formal analysis**: D.E., J.F.H., J.P.

**Investigation**: J.P

**Methodology**: D.E., J.P

**Project administration**: D.E.

**Software**: J.P.

**Supervision**: D.E.

**Validation**: D.E., J.P

**Visualization**: D.E., J.P

**Writing—original draft**: D.E., J.P

**Writing—review and editing**: D.E., J.F.H., J.P.

## A Appendix

### A.1 Statistical comparison of *G*_2_ and *G*_3_ against baseline

The following table reports the detailed test statistics along with the median score, measured using balanced accuracy for all classification tasks and r2-score for the age-regression.

**Table S1:** Detailed report of the test statistic.

| task | dataset | model | median score | t<sub>99</sub> | p-value |
| --- | --- | --- | --- | --- | --- |
| age prediction | TUAB | $G_3$ | 0.7211 | 2.9906 | 0.0018 |
| | | $G_2$ | 0.6879 | 0.8650 | 0.1946 |
| pathology classification | TUAB | $G_3$ | 0.8379 | 1.7064 | 0.0455 |
| | | $G_2$ | 0.8344 | 2.4251 | 0.0086 |
| EO vs EC | TDBRAIN | $G_3$ | 0.8623 | 2.2305 | 0.01400 |
| | | $G_2$ | 0.8223 | 0.2184 | 0.4138 |
| sex prediction | TDBRAIN | $G_3$ | 0.8325 | -0.0491 | 0.4804 |
| | | $G_2$ | 0.8141 | 1.9805 | 0.0252 |
| dementia diagnosis | CAU | $G_3$ | 0.6045 | 3.238332 | 0.0008 |
| | | $G_2$ | 0.5893 | 1.5949 | 0.0570 |

### A.2 Manifold optimization

Given a SPD input matrix, an unconstrained BiMap layer will output a symmetric positive **semi**-definite matrix, which means it can have eigenvalues that are zero. To ensure that the output of the BiMap layer is a SPD matrix, with strictly positive eigenvalues, the weights of the BiMap can be constrained to be semi-orthogonal. To do so, we used the trivialization for gradient-based optimization on manifolds introduced by Lezcano-Casado 2019. This method consists in parametrizing a manifold in terms of a Euclidean space. In *Theorem 4.3*, the author shows that this approach is theoretically equivalent to using Riemannian gradients (Absil et al., 2008) which has been used in previous publications using Riemannian geometry and DL for EEG (Carrara et al., 2024; Wilson et al., 2022). In addition to this theoretical result, we empirically observed that both approaches produce similar results. Given the improvement in computation time and reduced overhead, we opted for the trivialization approach.

### A.3 Visualization of wavelet transformed signals

**Figure S1:**
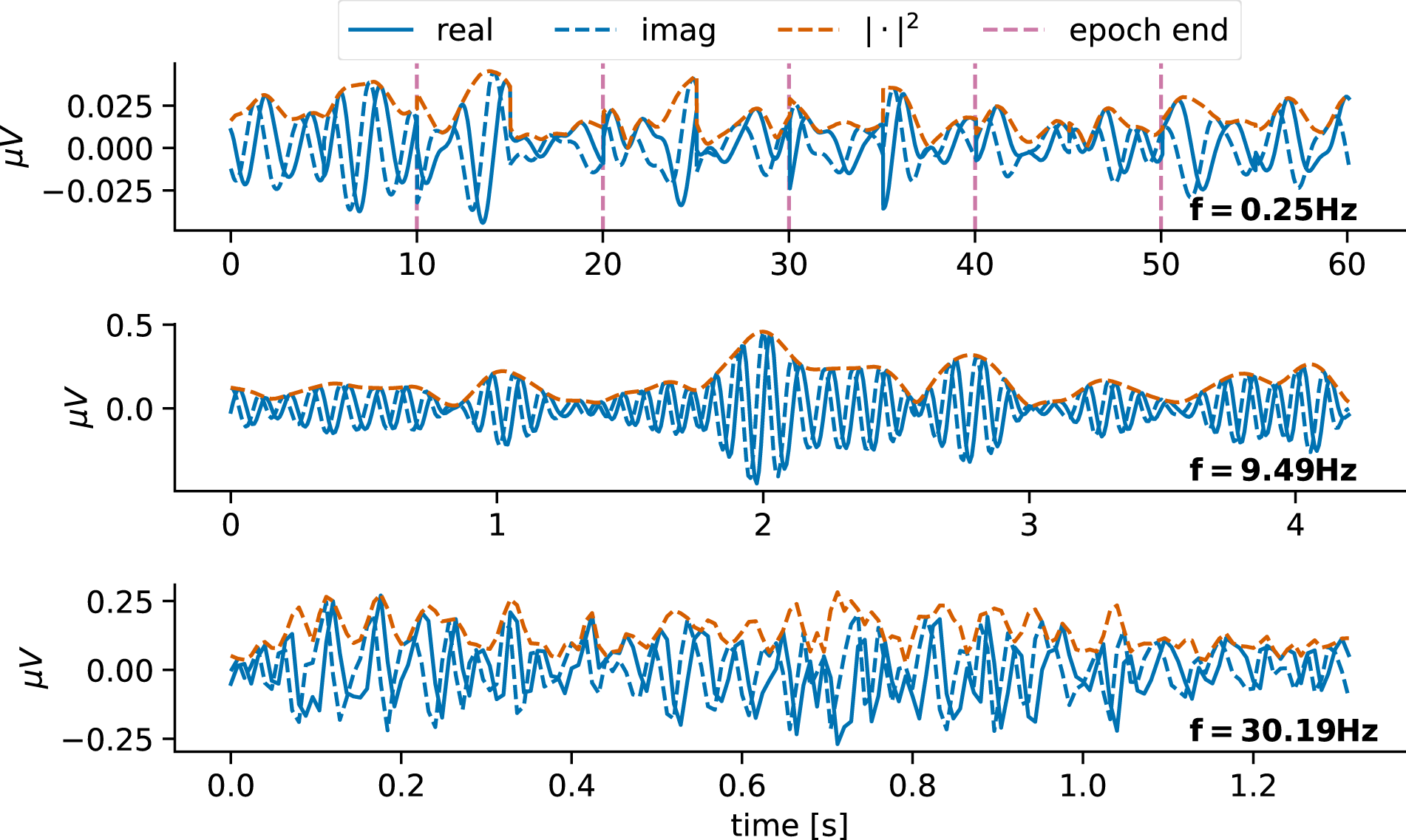
Wavelet-transformed signal. Each row represents the output of the convolution by one of the three complex-valued wavelets presented in Figure 4. The real and the imaginary parts are represented along with the squared modulus, sometimes referred to as power envelope. For the smallest frequency (0.25*Hz*), multiple epochs have been represented side by side. This plot also reveals the small variations captured by the wavelet, despite the fact that the signal has been high-pass filtered above 1*Hz*. This also suggests that task-relevant information is contained around this frequency, therefore pointing a sub-optimal preprocessing choice. The squared modulus variations are slower, evidencing its limited sensitivity to phase variations and small shifts.

## Footnotes

1 https://github.com/Roche/neuro-green

